# Substantial intraspecific variation in energy budgets: biology or artefact?

**DOI:** 10.1101/2021.01.07.425747

**Authors:** Tomos Potter, David N Reznick, Tim Coulson

## Abstract

1. Dynamic energy budget (DEB) models provide a mechanistic description of life-histories in terms of fluxes of energy through biological processes. In these models, life-histories are a function of environmental conditions and of fundamental traits of the organism relating to the acquisition, allocation, and use of energy.
2. These traits are described by the parameters of the DEB model, which have been estimated for over 2500 species. Recent work has aimed to compare species on the basis of differences in DEB parameters.
3. We show that caution is required in such analyses, because (i) parameter estimates vary considerably as an artefact of the types of data used to fit the models, and (ii) there is substantial intraspecific variation in parameter values, reflecting biological differences among populations.
4. We show that similar patterns of growth and reproduction can be reproduced with very different parameter sets. Our results imply that direct comparison of DEB parameters across populations or species may be invalid. However, valid comparisons are possible if differences in the types of data used to fit the models are taken into account.
5. We estimated DEB parameters for 16 populations of Trinidadian guppy, identifying differences in resource allocation and metabolic rate consistent with evolved life-history differences among these populations.
6. Variation in parameter values was substantial: if intraspecific variation in DEB parameters is greater than currently measured levels of interspecific variation, the detection of broad-scale patterns in energy budgets across species will be challenging.

## Introduction

The variety of life-history strategies observed in the natural world is truly astounding. From bristlecone pine that live for millenia, to mayflies that exist as adults for less than a day, evolution by natural selection has generated a vast array of solutions to life’s central problem: how best to convert energy into reproductive success. Classical theory proposes that life-history evolution is driven by the interaction between (i) extrinsic ecological effects on reproduction and survival; and (ii) intrinsic organismal constraints, arising from trade-offs in the allocation of energy to different biological processes (Stearns, 1989, 2000). Detection of these life-history trade-offs has been successful across taxa (Stearns, 1983; Bauwens and Díaz-Uriarte, 1997; Salguero-Gómez et al., 2016; Healy et al., 2019), but rarely so within species (van Noordwijk and de Jong, 1986). This is problematic, because intraspecific variation in (ii) is a prerequisite for life-history evolution to occur. Furthermore, life-history theory lacks specification of “the intra-organismal features that constrain the evolution of phenotypic traits”, limiting the potential for quantitative predictions of life-history evolution (Stearns, 2013). For a mechanistic understanding of life-history evolution, a model that captures the intrinsic trade-offs involved in developing and expressing traits that underpin the acquisition, allocation, and use of resources is required.

Dynamic Energy Budget (DEB) theory offers such a model (Kooijman, 2010). The standard DEB model describes individual life-histories, derived from first principles in accordance with the constraints of thermodynamic laws (Nisbet et al., 2000; Sousa et al., 2008; Jusup et al., 2017). The model provides a single framework to describe an animal’s development, growth, maintenance, reproduction, and ageing, under any range of environmental conditions (Kooijman, 2010). Models are parameterised by quantifying the rates of assimilation, allocation, and efficiency of energy flow between biological processes under varying levels of food availability and/or temperature (van der Meer, 2006; Kooijman et al., 2008; Kooijman, 2010). The primary parameters of the standard DEB model represent fundamental (albeit abstract) traits of individuals, that, along with resource availability and temperature, determine the life-history of the organism (Kooijman, 2010). DEB models can be parameterised with a wide range of commonly collected empirical data, and to date parameters have been estimated for over 2500 species, covering all chordate classes (Marques et al., 2018; Augustine et al., 2019; AmP, 2020).

Differences in DEB parameter values between species are assumed to reflect evolved differences (Kooijman, 2010; Lika et al., 2011a; Marques et al., 2018; Lika et al., 2020). The reasoning underpinning this assumption is that the structure of the DEB model means that parameters should co-vary in a simple way with maximum body size: differences in parameter values between species (accounting for differences in maximum body size) therefore indicate departures from this “null model” and imply evolutionary adaptations of metabolic processes (Kooijman, 2010; Lika et al., 2011a; Marques et al., 2018). Recent work has begun to explore patterns of parameter variation across species, identifying several spectra of life-history strategies which are defined by the different ways in which species allocate energy to development, growth, and reproduction (Marques et al., 2018; Lika et al., 2019; Augustine et al., 2019).

Although theory defines DEB parameters as being individual-specific, individual-level differences are considered to be sufficiently small that mean values are taken to represent species-specific parameters (Marques et al., 2018). This assumption is central to analyses of interspecific patterns of variation in DEB parameters, yet little is known about how much these parameters actually vary within species. To our knowledge, only two studies have employed DEB theory to quantify life-history differences between natural populations of the same species (Marn et al., 2019; Guillaumot et al., 2020). Marn and colleagues (2019) found that DEB parameters differed between Mediterranean and North Atlantic loggerhead turtles, and that life-history differences between these populations could not be explained as a direct result of environmental differences between their respective habitats. In contrast, Guillaumot and colleagues (2020) were unable to determine differences in DEB parameters between intertidal and subtidal ecotypes of the Antarctic limpet, despite clear morphological divergence. As such, there is little consensus on the extent to which DEB parameters can vary within a single species.

Here, we assess the extent of intraspecific variation in energy budgets of the Trinidadian guppy (*Poecilia reticulata*). Guppies display well characterised variation in life-history strategies, evolving distinct ecotypes in the presence or absence of predators (Reznick and Endler, 1982; Reznick, 1982; Reznick and Bryga, 1987). Ancestral populations of guppies live in habitats characterised by the presence of voracious predators, in particular the pike cichlid (*Crenicichla alta*) (Magurran, 2005). In several rivers, guppies have invaded further upstream, beyond barrier waterfalls which restrict the movement of large predatory fish. In these upstream habitats, guppies coexist with only one other fish species, Hart’s killifish (*Anablepsoides hartii*), which is a competitor, and also an occasional predator of guppies (Magurran, 2005). On release from intense predation pressure, these newly founded upstream guppy populations grew rapidly, until diminishing resource availability regulates population growth, driving selection for a slower life-history (Bassar et al., 2013; Reznick et al., 2019; Potter et al., 2021). The “low-predation” ecotype matures at larger sizes and older ages, and produces fewer, larger offspring per litter than their “high-predation” ecotype ancestors (Reznick and Endler, 1982; Reznick, 1982; Reznick and Bryga, 1987). The low-predation ecotype has repeatedly and independently evolved several times in Trinidad, both in natural and experimentally introduced populations (Reznick, 1982; Shaw et al., 1992; Alexander et al., 2006; Reznick et al., 2019; Whiting et al., 2020). As such, Trinidadian guppies provide an excellent system in which to test the ability of DEB theory to distinguish intraspecific variation in the intrinsic trade-offs that underpin life-history evolution.

In this study, we investigate the degree of variation in DEB parameters among 16 populations of Trinidadian guppies, and assess whether DEB theory can predict known life-history differences among ecotypes of this species. We find evidence for substantial variation in energy budgets, reflecting both biological differences between populations, and artefacts of the parameter estimation process. We discuss our findings in the context of quantifying within-species variation in the intrinsic trade-offs that govern life-histories.

## Methods

### Overview of methods

We estimated the parameters of the standard dynamic energy budget model for sixteen guppy populations, covering five independent evolutionary origins of the low-predation ecotype. According to DEB theory, differences in parameter values between populations indicate evolved differences in fundamental processes underlying the acquisition, allocation and use of energy. However, differences in parameter values may also arise from differences in the types of data used to fit the models (Lika et al., 2011b). To assess this effect, we fit DEB models for four focal populations with three different levels of data availability. To identify DEB parameters associated with adaptation to resource limitation, we quantified differences in parameters between guppy ecotypes, assessing the generality of these differences across all populations fit with the same degree of data availability. Finally, we compared the variance in parameter values within Trinidadian guppies with that reported across species within the order Cyprinodontiformes.

### The standard DEB model

The standard dynamic energy budget model (hereafter ‘DEB model’) describes how an organism acquires, allocates, and uses energy over the course of its life, from its beginning as a fertilised egg until its death through ageing (Kooijman, 2001; Sousa et al., 2008; Kooijman, 2010). For a detailed description and derivations of the full model we refer the reader to Chapter 2 of Kooijman (2010). The DEB model defines three distinct life-history stages: embryonic (no eating; no reproduction), juvenile (eating; no reproduction), and adult (eating; reproducing). An individual is characterised by four state variables: reserve, structure, maturity level, and reproductive buffer (Fig. 1). Reserve can be considered as stored energy, obtained through feeding, which fuels metabolic processes. Structure describes the soma of the organism, and can be considered as the physical structure of the body produced and maintained through the metabolization of reserve. The transitions between life-history stages (i.e. embryonic to juvenile; juvenile to adult) are determined by maturity level. This somewhat abstract concept captures the cumulative energy spent in developing physiological complexity, including development of the immune system (Kooijman, 2010). The reproductive buffer describes the amount of mobilised reserve that is allocated specifically to reproduction. These states vary over the life of the organism as a function of the parameters of the model (Table 1) and their interactions with environmental conditions (namely, temperature and food availability). The model captures the biological processes of feeding, assimilation of food, defecation, mobilisation of assimilated energy, allocation of mobilised energy to somatic or reproductive work, maintenance costs of somatic and reproductive work, somatic growth, attainment of maturity, reproduction, and ageing (Figure 1, Table 1, Table 2). Data on the growth and reproduction of individuals at more than one level of food availability are considered sufficient for estimation of most of the primary parameters of the DEB model (Lika et al., 2011a).

**Table 1:** Parameters of the standard DEB model and the primary biological processes that they control. Two further parameters, the shape coefficient δ_*m*_ and the zoom factor *z*, fully describe the shape of the organism, link structure *V* to physical measurements, and allow comparisons between species of different size and shape. Symbols follow the standard DEB notation (Kooijman, 2010).

| Symbol | Description | Units | Process |
| --- | --- | --- | --- |
| $\{\dot{F}_m\}$ | Maximum specific searching rate | $\text{L d}^{-1} \text{ cm}^{-2}$ | Feeding |
| $\{\dot{p}_{Am}\}$ | Maximum specific assimilation rate | $\text{J d}^{-1} \text{ cm}^{-2}$ | Assimilation |
| $\kappa_X$ | Efficiency of converting food to reserve | - | Defecation |
| $\dot{v}$ | Energy conductance | $\text{cm d}^{-1}$ | Mobilisation of reserve |
| $\kappa$ | Fraction of reserve allocated to somatic work | - | Allocation of mobilised reserve |
| $\kappa_R$ | Reproductive efficiency (reserve to egg) | - | Reproduction |
| $[\dot{p}_M]$ | Volume-specific somatic maintenance cost | $\text{J d}^{-1} \text{ cm}^{-3}$ | Somatic maintenance |
| $\{\dot{p}_T\}$ | Surface-area specific somatic maintenance cost | $\text{J d}^{-1} \text{ cm}^{-2}$ | Somatic maintenance |
| $k_{\mathcal{J}}$ | Maturity maintenance rate coefficient | $\text{d}^{-1}$ | Maturity maintenance |
| $[E_G]$ | Specific cost of structure | $\text{J cm}^{-3}$ | Somatic growth |
| $E_H^b$ | Maturity at birth | J | Life-history: onset of feeding |
| $E_H^p$ | Maturity at puberty | J | Life-history: onset of allocation to reproduction |
| $\dot{h}_a$ | Weibull ageing acceleration | $\text{d}^{-2}$ | Ageing |
| $S_G$ | Gompertz stress coefficient | - | Ageing |

**Table 2:**
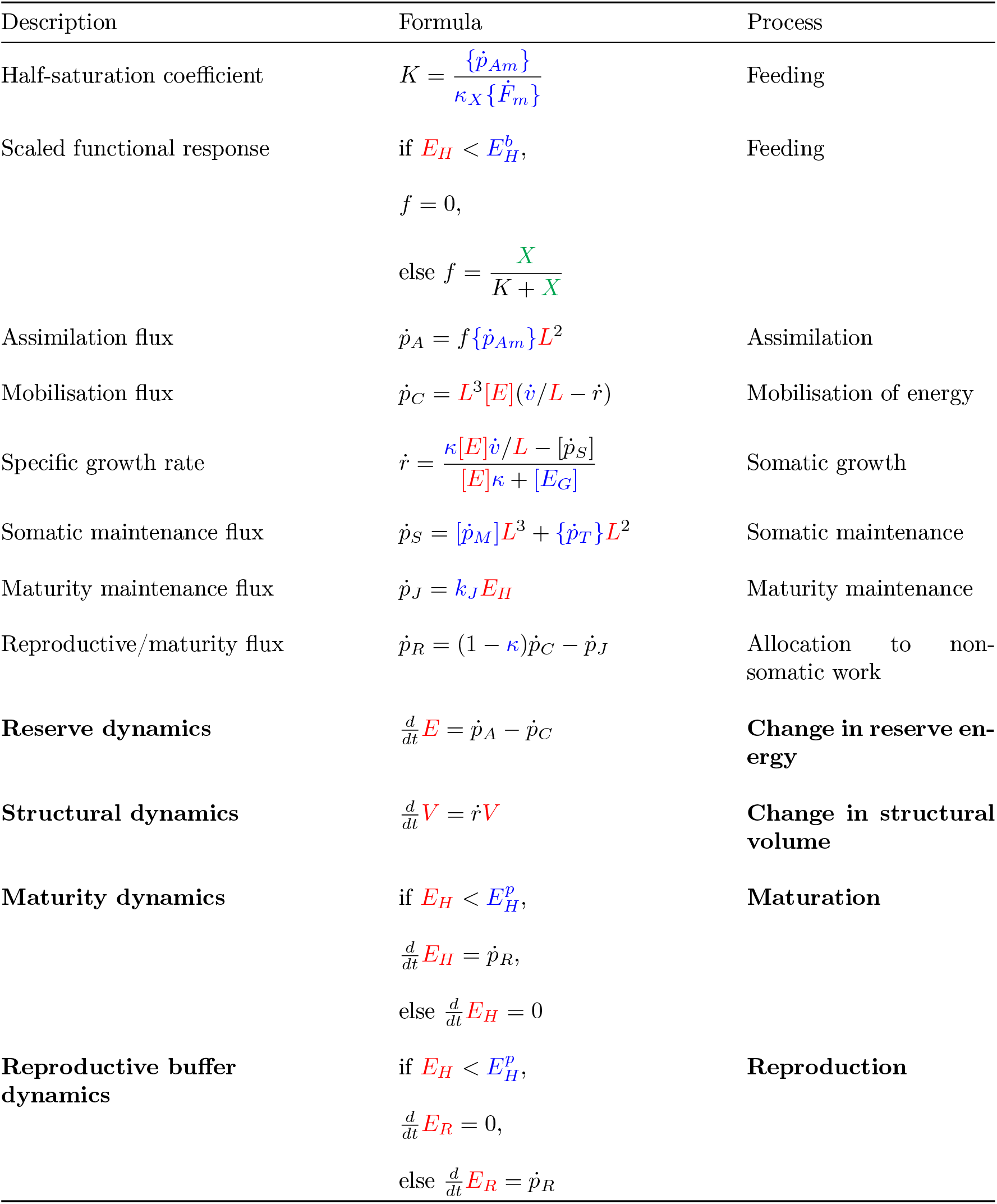
Linking primary parameters to biological processes in the standard DEB model. Formulae are summarised from Table 1 in Lika & Kooijman (2011). The state variables (in red) are structural volume *V* (cm^3^), energy in the reserve *E* (J), energy allocated to maturity *E_H_* (J), and energy available for reproduction *E_R_* (J). To simplify the presentation of equations, we also refer to two “compound” state variables: structural length *L* (*L* = *V* ^1/3^, cm), and reserve energy density [*E*] ([*E*] = *E/V*, J cm^−3^). Primary parameters (in blue) are described in Table 1. Environmental food density *X* is given in green. Compound parameters and state functions (in black) are defined within this table, and are functions of state variables, primary parameters, and food density. Symbols follow the standard DEB notation (Kooijman, 2010). Values that are expressed per unit of structural volume are in square brackets [], those expressed per unit of structural surface area are in braces {}.

**Figure 1:**
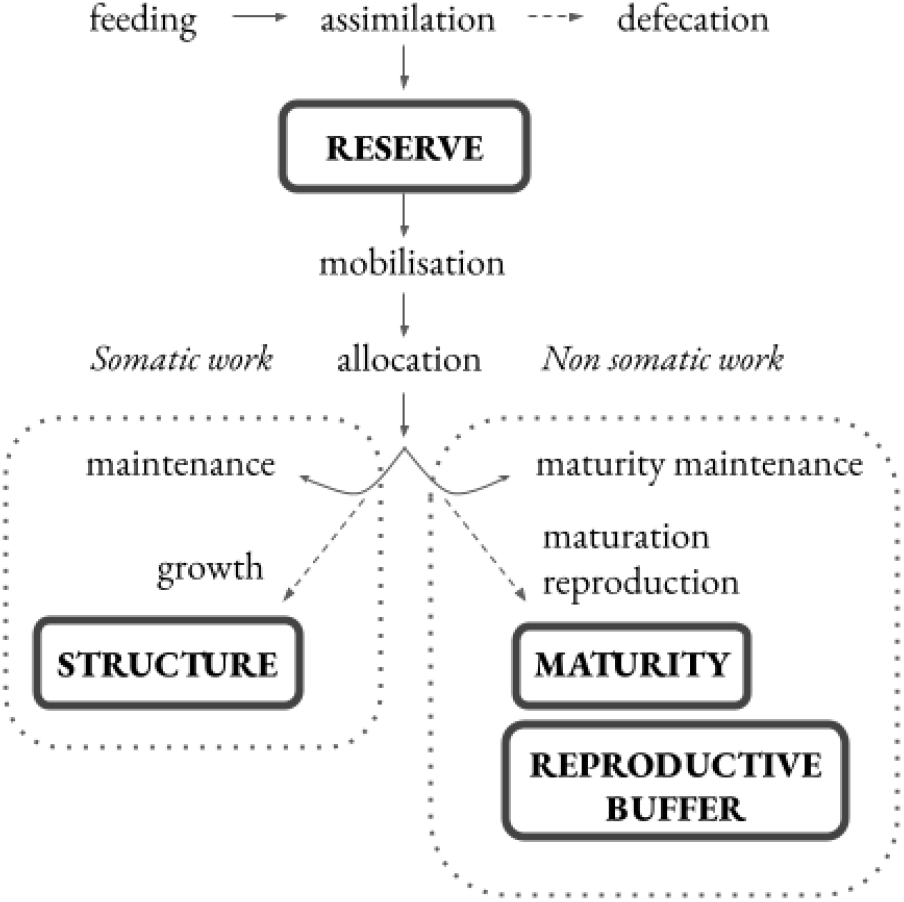
Schematic of the standard dynamic energy budget model. State variables are given in boxes, processes are shown in plain text. Processes (including ageing, not depicted) are determined by 14 parameters, described in Table 1. Arrows represent fluxes of energy through the system via biological processes and state variables. A fraction *κ* of mobilised reserve is allocated to somatic work, where somatic maintenance costs must first be met before the remainder is allocated to growth. Reserve allocated to reproductive work (1 − *κ*) is first used to meet maturity maintenance costs, and then used to attain sexual maturity (in juveniles) or produce offspring (in adults).

### Estimating DEB parameters and predicting metabolic rate

The majority of the data that we used to fit population-specific DEB models came from three sets of common garden experiments, in which guppies sampled from sixteen population were reared at either high or low levels of food availability (Reznick and Bryga, 1996; Reznick et al., 2004, 2005, and one unpublished dataset from D. Reznick and colleagues). Details of how these data were collected, including information on the sampled populations (location, sample size, sampling year etc), rearing protocols, and trait measurements, are provided in the supplementary material.

Data used to fit DEB models are of two types: zero-variate, and uni-variate (Lika et al., 2011b). Zero-variate data are single values, whereas uni-variate data captures the relationship between pairs of values.

For our study, zero-variate data included the age at birth (mean inter-birth interval, in days) and the mean dry weight of offspring (g), with both measures recorded at high and low food levels, and ultimate standard length (cm) at *ad lib* food. Univariate data included age (days since birth) versus standard length (cm), weight (g), cumulative reproductive output (number of offspring), and length versus weight. Each of these measures were recorded at high and low food levels. For twelve of our study populations, these data were recorded at the first three parturition events only. For four populations, data were recorded at each parturition event over the entire lifespan of individuals. In addition, for these four populations we recorded mean dry weight (mg) of neonates at high and low food, the surviving fraction of the population as a function of time since birth, daily food ration (joules per day) used in the common garden experiments (Reznick et al., 2004), and we also incorporated data on basal metabolic rate (oxygen consumption at 25 °C, mg hr^−1^) from an independent study using the same populations (Auer et al., 2018).

For four of our focal populations, more types of data were available to estimate DEB parameters than in the other twelve. To assess how the use of different types of data may affect the estimation of parameters (Lika et al., 2011a), we first fit DEB models for the first four populations at three different levels of data availability (Table 3). To compare populations across all datasets, we then fit sixteen population-specific DEB models using the same level of data availability (Table 3).

**Table 3:** Types of data used to estimate DEB parameters in this study at each level of data availability tested. For twelve populations, we used data at availability level 1 only. For four populations, data was available at all three levels. The asterisk * indicates that level 1 univariate data were available for measurements at the first three parturition events only, i.e. 3 observation per individual. Univariate data at levels 2 and 3 were recorded at each parturition event over the full lifespan: mean number of observations per individual=22 (s.d.=8.5).

| Data type | Units | Food treatment | Data level |
| --- | --- | --- | --- |
| Age at birth | d | high, low | 1,2,3 |
| Maximum standard length | cm | ad lib | 1,2,3 |
| Age vs weight | d, g | high, low | 1*,2,3 |
| Age vs length | d, cm | high, low | 1*,2,3 |
| Length vs weight | cm, g | high, low | 1*,2,3 |
| Age vs cumulative reproduction | d, - | high, low | 1*,2,3 |
| Dry weight at birth | mg | high, low | 2,3 |
| Age vs surviving fraction of population | d, - | high, low | 2,3 |
| Age vs food availability | d, J d<sup>-1</sup> | high, low | 3 |
| Dioxygen consumption rate | mg hr<sup>-1</sup> | ad lib | 3 |

Both zero-variate and uni-variate observations can be described as functions of primary DEB parameters, state variables, temperature, and food availability. Because DEB parameters appear in multiple auxiliary models of observable relationships, model fitting is not a straight-forward process. All relationships must be modelled simultaneously because no single parameter is specific to a single type of observation (Marques et al., 2019). DEB parameters are derived from theory, meaning that there are substantial constraints on their values. For example, *κ*, the fraction of energy allocated to somatic work, must be bounded between 0 and 1, and the energy conductance rate 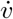 must be positive, etc. This means that parameter filters and ‘psuedo-data’ – parameter values that correspond to a generalised animal where the zoom factor *z*=1 – can be used to constrain the parameter space during estimation, in a manner that is analogous to the use of priors in Bayesian estimation (Lika et al., 2011a, 2014; Marques et al., 2019). The use of pseudo-data increases the identifiability of parameters, whilst their influence on final parameter estimates is restricted by low weighting of pseudo-data during estimation. Parameter estimation is based on the minimisation of a symmetric-bounded loss function, using a Nealder-Mead simplex minimization approach (Marques et al., 2019).

We estimated parameters from data using the freely available software package DEBtool_M (available from github.com/add-my-pet/DEBtool_M, (Marques et al., 2018)) in MATLAB version R2020a. We used standard functions in the DEBtool_M package to predict the data for analyses with data availability levels 1 and 2. For analyses including data on food availability we built a custom function, consisting of a set of differential equations which described the rates of change of the reserve, structure, maturity level, reproductive buffer, and survival probability, accounting for the changes in food availability with age. We linked the changes in state variables to observations of traits via standard auxiliary theory (Kooijman, 2010). Biological rates are temperature dependent: for all analyses, we assumed an Arhennius temperature of 8000 K. Some DEB parameters could not be estimated from the available data: we fixed the maximum specific searching rate 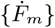 at 6.5 L d^−1^ cm^−2^, and the reproductive efficiency *κ_R_* at 0.95 in all analyses – these are typical parameter values, and are recommended for use when no data is available for estimation (Lika et al., 2011a). Without information on food availability (i.e. data levels 1 and 2), assimilation efficiency *κ_X_* cannot be estimated: we fixed this value at 0.25 based on our estimates of *κ_X_* at data availability level 3. When we could not directly estimate *κ_X_* we quantified feeding rates by estimating the scaled functional response *f* (Table 2) at high and low food levels. To achieve a better fit of the models to the data, we introduced two additional parameters: the first captured a delay in the onset of development, *T_0_* (measured in days) and the second was a low-food level specific assimilation efficiency 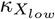. The parameter *T_0_* was included as an additive term in the auxiliary model for age at birth only. For modelling growth, reproduction, and survival at low food levels, we used 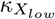 in place of *κ_X_* in the standard DEB model.

Although metabolic rate data (oxygen consumption) were only used to fit models with data availability level 3 (Table 3), we predicted metabolic rates from the parameter sets for each population, using the “scaled_power” function in DEBtool_M. For these predictions, we estimated the total power (i.e. energetic flux *J d* ^−1^) expended by a guppy weighing 74 mg, at a temperature of 25 °C. We excluded assimilation power from our analyses, because our reference data were collected from guppies that had fasted for 24 hours before measurement (Auer et al., 2018). We then converted this energetic flux into a mass flux (milligrams of O_2_ per hour) following standard assumptions about the chemical composition and potential (J mol^−1^) of structure and reserve, and assuming a molar weight for dioxygen of 32 g mol^−1^ (Kooijman 2010, Chapter 3).

Each population-specific set of data and the MATLAB scripts required to estimate the parameters presented in this manuscript are available to download from the AmP online repository (AmP, 2020) (see Table S1).

### Assessing model fit and differences among parameter sets

We assessed goodness of fit of the models to the data by calculating the mean error of the predictions relative to the data, for each set of observations used to fit the model (Marques et al., 2018). The overall fit of the model is described by the mean relative error, which is the grand mean of the relative error terms from each set of predictions and observations.

To determine whether there were significant differences in parameter sets between populations, we followed the approach described by Marn and collegaues (2019): we compared the fit of the model to the data when using the population-specific (“best”) parameters with that when using a null-model parameter set. We compared the “best” parameters with two null models: “null 1”, which was the parameter set of the alternative ecotype from the same stream (e.g. LP Yarra vs HP Yarra); and “null 2”, which was the parameter set of the same ecotype from the alternative stream (e.g. LP Yarra vs LP Oropuche). This allowed us to test whether the parameter sets were distinct between (i) ancestral and derived populations in the same stream, and (ii) between common ecotypes that have evolved in different streams. We assessed differences in parameter sets between the four populations (two ecotype pairs) for which the most data were available to fit the models. We tested the difference in fit of DEB parameter sets using Wilcoxon’s paired rank test (Marn et al., 2019).

Finally, we assessed how intraspecific variation in DEB parameters compares with interspecific variation. Parameter values are expected to be more similar among closely related species. We compared parameter values obtained in this study for Trinidadian guppies with parameter values of other species within the order Cyprinodontiformes. We retrieved all available Cyprinodontiformes parameter sets from the AmP database (AmP, 2020) (59 species), and visualised the variation among pairs of parameters. We performed F-tests to compare intra- and interspecific variation in DEB parameters. Calculations were performed in the in the R environment (R Core Team, 2019), and all figures were produced using the “ggplot2” package (Wickham, 2011).

## Results

### Impact of data availability on parameter estimates

The estimation of dynamic energy budget model parameters depended on the type and amount of data used to fit the models (Table 4). Estimates of the fraction of energy allocated to somatic work *κ*, the maximum assimilation rate 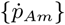, and of the volume-specific somatic maintenance rate 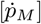 were all substantially lower when estimated using data level 3. Estimates of the maturity threshold at the onset of the adult stage, 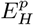, were highest when estimated using level 3 data. These patterns were consistent across all four populations tested (Table 4).

**Table 4:** Estimates of dynamic energy budget model parameters vary depending on the type and amount of data used to fit the models. We estimated parameters using three levels of data availability (see Table 3). Parameters are shown for four populations of Trinidadian guppy: the high-predation (HP) and low-predation (LP) ecotypes from the Oropuche and Yarra rivers. Parameters which were fixed and therefore not estimated during model fitting are denoted with (n. e.). Parameters that were fixed across all data levels were: the maximum specific searching rate 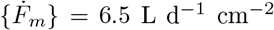; the reproductive efficiency *κ_R_* = 0.95; the surface area specific somatic maintenance 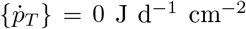; and the Arhennius temperature = 8000 K. Models were fit assuming an environmental temperature of 26 °C; following convention, we report temperature-adjusted parameters for a reference temperature of 20 °C. The mean relative error (MRE) quantifies the fit of the models to the data.

| Parameter | Population:<br>Data level:<br>Unit | Oropuche HP |  |  | Oropuche LP |  |  | Yarra HP |  |  | Yarra LP |  |  |
| --- | --- | --- | --- | --- | --- | --- | --- | --- | --- | --- | --- | --- | --- |
|  |  | 1 | 2 | 3 | 1 | 2 | 3 | 1 | 2 | 3 | 1 | 2 | 3 |
| Assimilation efficiency, $\kappa_X$ | - | 0.25<sup>ne</sup> | 0.25<sup>ne</sup> | 0.27 | 0.25<sup>ne</sup> | 0.25<sup>ne</sup> | 0.23 | 0.25<sup>ne</sup> | 0.25<sup>ne</sup> | 0.26 | 0.25<sup>ne</sup> | 0.25<sup>ne</sup> | 0.21 |
| Assimilation efficiency at low food, $\kappa_X^{low}$ | - | - | - | 0.34 | - | - | 0.30 | - | - | 0.34 | - | - | 0.26 |
| Max specific assimilation rate, $\{\dot{p}_{Am}\}$ | $\text{J d}^{-1} \text{ cm}^{-2}$ | 116.7 | 117.8 | 63.1 | 108.2 | 132.1 | 52.5 | 175.0 | 138.1 | 74.6 | 156.4 | 110.9 | 56.1 |
| Energy conductance, $\dot{v}$ | $\text{cm d}^{-1}$ | 0.028 | 0.028 | 0.029 | 0.029 | 0.033 | 0.025 | 0.029 | 0.028 | 0.030 | 0.027 | 0.030 | 0.030 |
| Allocation fraction to soma, $\kappa$ | - | 0.55 | 0.73 | 0.43 | 0.70 | 0.78 | 0.58 | 0.61 | 0.70 | 0.40 | 0.69 | 0.78 | 0.54 |
| Vol-spec somatic maint, $[\dot{p}_M]$ | $\text{J d}^{-1} \text{ cm}^{-3}$ | 68.2 | 93.5 | 27.31 | 95.1 | 119.4 | 35.32 | 120.1 | 114.5 | 33.78 | 121 | 105.7 | 34.31 |
| Maturity maint rate coefficient, $\dot{k}_J$ | $\text{d}^{-1}$ | 0.0019 | 0.0020 | 0.0020 | 0.0020 | 0.002098 | 0.0017 | 0.0020 | 0.0020 | 0.0020 | 0.0020 | 0.0021 | 0.0021 |
| Spec cost for structure, $E_G$ | $\text{J cm}^{-3}$ | 5220 | 5220 | 5223 | 5228 | 5219 | 5185 | 5225 | 5221 | 5213 | 5222 | 5220 | 5232 |
| Maturity at birth, $E_H^b$ | $\text{J}$ | 36.9 | 6.4 | 26.6 | 12.8 | 6.7 | 17.9 | 25.2 | 5.5 | 21.6 | 8.8 | 5.3 | 18.1 |
| Maturity at puberty, $E_H^p$ | $\text{J}$ | 298.3 | 63.0 | 501.8 | 146.5 | 222.6 | 401.5 | 382.5 | 264.9 | 515.3 | 238.9 | 227.7 | 524.5 |
| Weibull aging acceleration, $\dot{h}_a$ | $\text{d}^{-2}$ | - | $2.65 \times 10^{-8}$ | $2.03 \times 10^{-8}$ | - | $3.98 \times 10^{-8}$ | $4.10 \times 10^{-8}$ | - | $6.15 \times 10^{-8}$ | $4.41 \times 10^{-8}$ | - | $6.71 \times 10^{-8}$ | $5.62 \times 10^{-8}$ |
| Gompertz stress coefficient, $S_G$ | - | - | 0.070 | 0.062 | - | 0.074 | 0.075 | - | 0.029 | 0.037 | - | 0.048 | 0.049 |
| Zoom factor, $z$ | - | 0.94 | 0.92 | 0.99 | 0.80 | 0.87 | 0.86 | 0.89 | 0.84 | 0.88 | 0.89 | 0.82 | 0.88 |
| Shape coefficient, $\delta_M$ | - | 0.24 | 0.24 | 0.26 | 0.24 | 0.25 | 0.26 | 0.23 | 0.23 | 0.25 | 0.23 | 0.24 | 0.26 |
| Scaled functional response high food, $f_{high}$ | - | 0.86<sup>ne</sup> | 0.92 | - | 0.86<sup>ne</sup> | 0.93 | - | 0.86<sup>ne</sup> | 0.90 | - | 0.86<sup>ne</sup> | 0.96 | - |
| scaled functional response low food, $f_{low}$ | - | 0.79 | 0.72 | - | 0.79 | 0.74 | - | 0.81 | 0.72 | - | 0.76 | 0.75 | - |
| Developmental delay, $t_0$ | $\text{d}$ | 5.2 | 8.171 | 6.7 | 9.0 | 10.7 | 7.7 | 6.0 | 8.149 | 7.0 | 9.8 | 10.32 | 8.9 |
| Model fit, MRE | - | 0.111 | 0.076 | 0.076 | 0.113 | 0.087 | 0.099 | 0.122 | 0.089 | 0.086 | 0.120 | 0.089 | 0.085 |

Despite substantial differences in parameter values by data level, the different parameter sets were able to reproduce how standard length, mass, and cumulative reproductive output changed as a function of age and food level (Fig. 2), as well as predicting age and dry weight at birth (at high and low food) and maximum length (Table S2). The overall fits of the models to the data, as quantified by the mean relative error, were poorest for models fit with data level 1, and were reasonable (MRE<0.1) and comparable between models fit at data levels 2 and 3 (Table 4).

**Figure 2:**
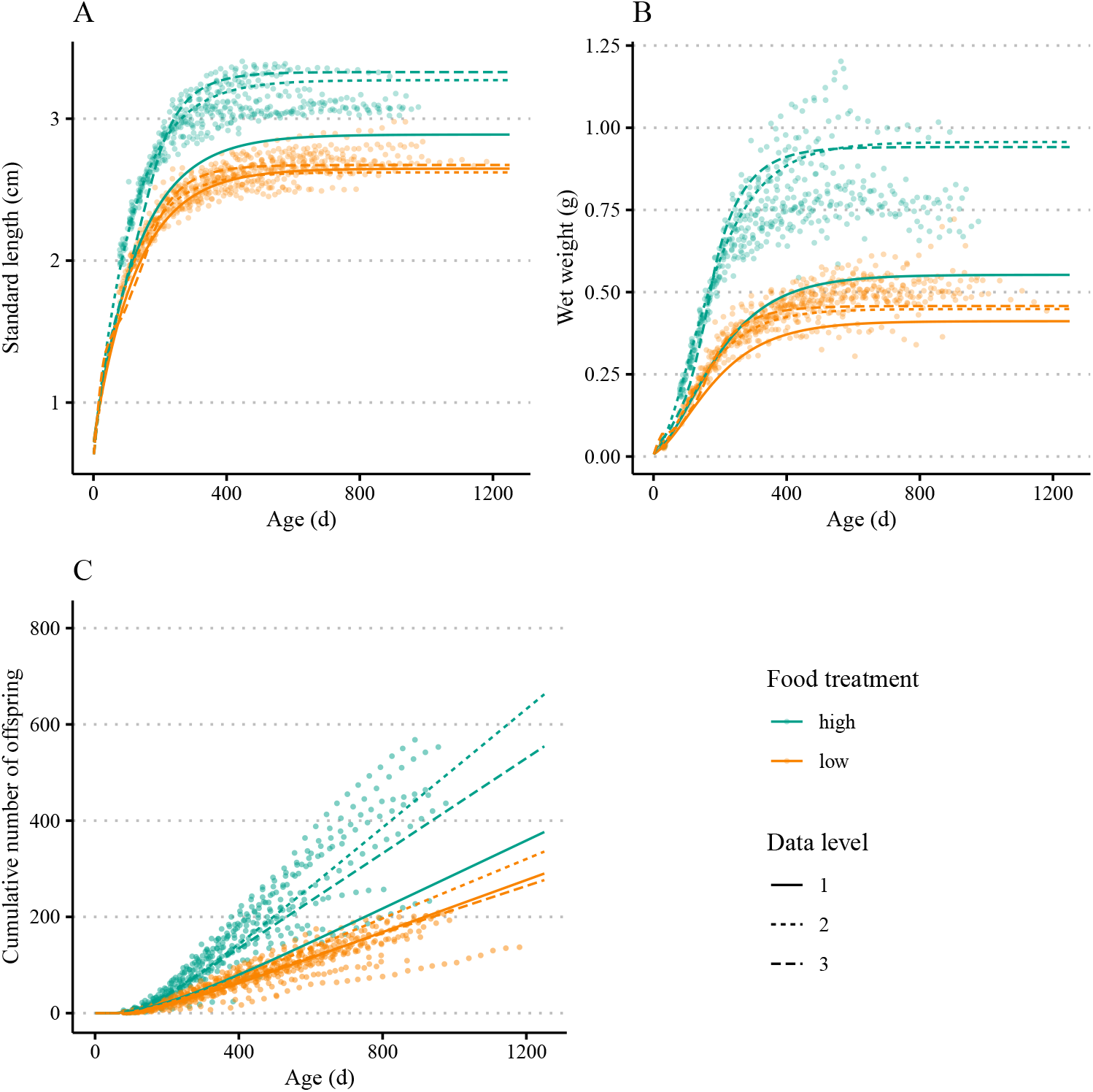
Observations and predictions of (A) length, (B) weight, and (C) cumulative number of offspring, as a function of age. Data (points) and predictions (lines) are of low-predation ecotype guppies from the Oropuche river, under high (blue) and low (orange) food treatments. Lines show predictions based on different parameter sets, estimated with different levels of data availability (defined in Table 3).

### Ecotype-specific parameter differences

Our initial assessment of ecotype-specific differences in DEB parameters considered two ecotype pairs (four populations) for which level 3 data were available. In general, parameter sets were significantly different between populations and ecotypes: the population-specific parameter sets gave a significantly better fit to the data compared to parameter sets from the other population (different ecotype) within the same river systems, or the from the same ecotype from a different river system (Table S3). There was one exception: for data from the low-predation Oropuche population, there was no significant difference in fit when using the population-specific parameter set, or the parameter set from the low-predation Yarra population (Table S3).

Guppy ecotypes adapted to high and low predation-risk environments differ in how they obtain, allocate, and use resources: we found that DEB parameters differed between ecotype pairs, and that these differences were often consistent between independent evolutionary origins of ecotype pairs in the Oropuche river and Yarra rivers. (Table 4, Fig. 3). Six parameters varied consistently between ecotypes: for both rivers, compared to the high-predation ecotype, low-predation guppies had lower maximum assimilation rates 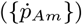 and assimilation efficiencies (*κ_X_* and *κ_X low_*), allocated a greater fraction of available energy to somatic work (*κ*), had a longer delay period at the onset of development (*t_0_*), and had higher values for the Gompertz stress coefficient (*S_G_*) (Fig. 3A).

**Figure 3:**
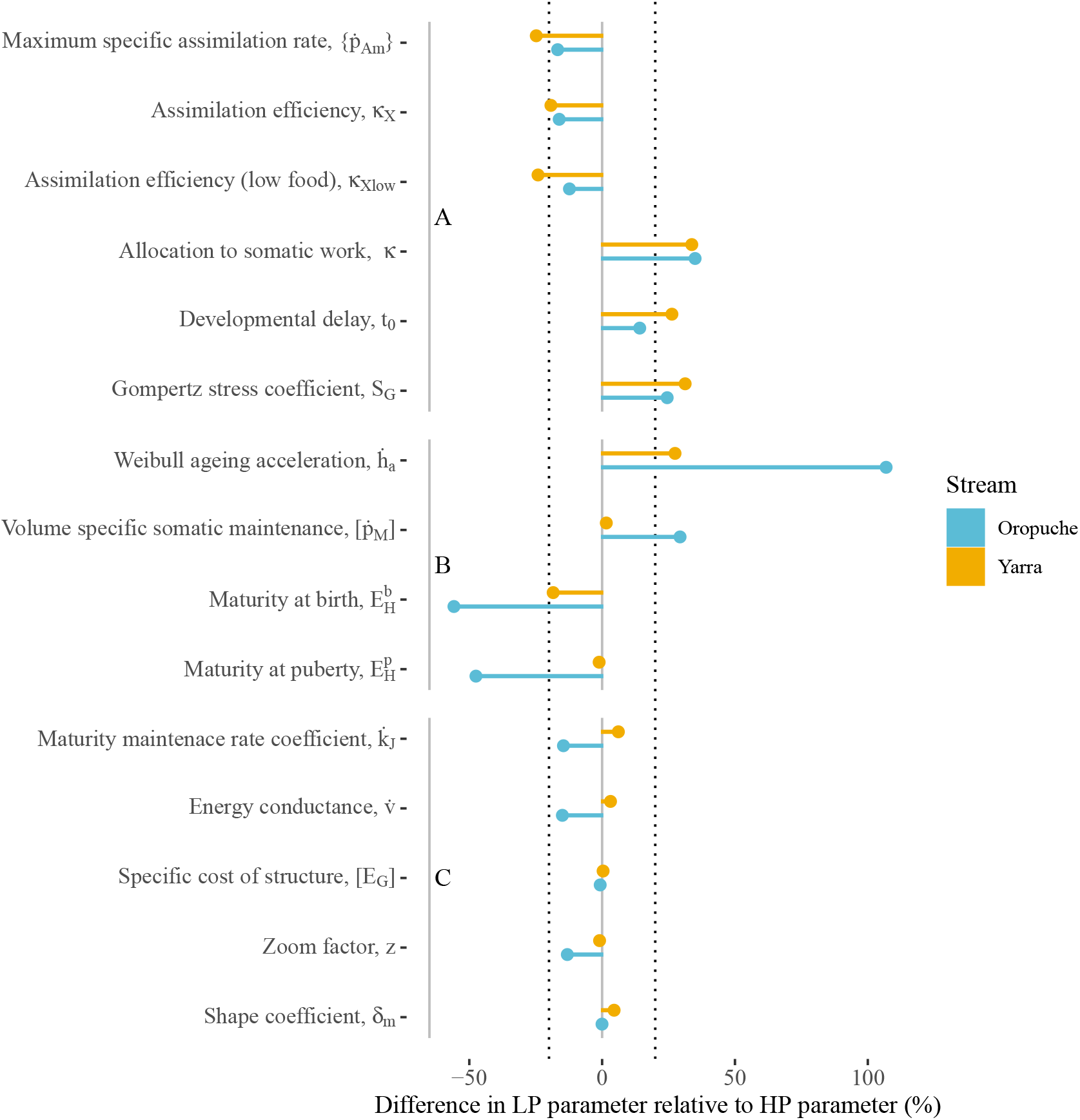
Percentage differences in DEB parameter values of low-predation (LP) relative to high-predation (HP). Differences between ecotypes from the Oropuche are given in blue; those from the Yarra are given in yellow. Parameters are categorised into three groups: (A) Parameters differ between ecotypes, and the direction of the effect is consistent between streams; (B) Parameters differ between ecotypes, and the direction and/or magnitude of the effect differs between streams; and (C) Parameters do not differ between ecotypes or streams. Dotted lines indicate a difference of plus or minus 20%, representing the threshold used to categorise parameters as falling in A, B, or C.

The magnitude of ecotype differences were inconsistent between streams for somatic maintenance costs 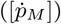, the Weibull ageing acceleration parameter 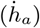, and for maturity thresholds (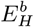 and 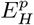) (Fig. 3B).

For five parameters, there were negligible differences between ecotypes, and this was consistent between streams. These parameters were the maturity maintenance coefficient 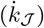, energy conductance 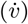, the energetic cost of structure ([*E_G_*]) the shape coefficient (δ_*m*_), and the zoom factor *z* (Fig. 3C).

We then fit DEB models at level 1 data availability for these four populations, plus an additional twelve populations. The overall fits of the models (MRE) ranged from 0.076 to 0.137, with a grand mean MRE of 0.107. Full parameter sets and model fit scores are provided in Table S4. We again found that the low-predation ecotype allocated a greater fraction of energy to somatic work, and had lower maximum assimilation rates than the high-predation ecotype (Fig. 4). Using the parameter sets for each population, we predicted basal metabolic rate in terms of dioxygen consumption rates. Note that metabolic rate data was not used to fit models at data level 1. The low-predation ecotype was predicted to have lower rates of oxygen consumption than the high-predation ecotype (Fig. 4C) – a pattern consistent with previously published work (Auer et al., 2018). However, the estimates of oxygen consumption for both ecotypes were typically about 50% greater than the values of those reported in (Auer et al., 2018), i.e. the values used to fit models at level 3 in this study.

**Figure 4:**
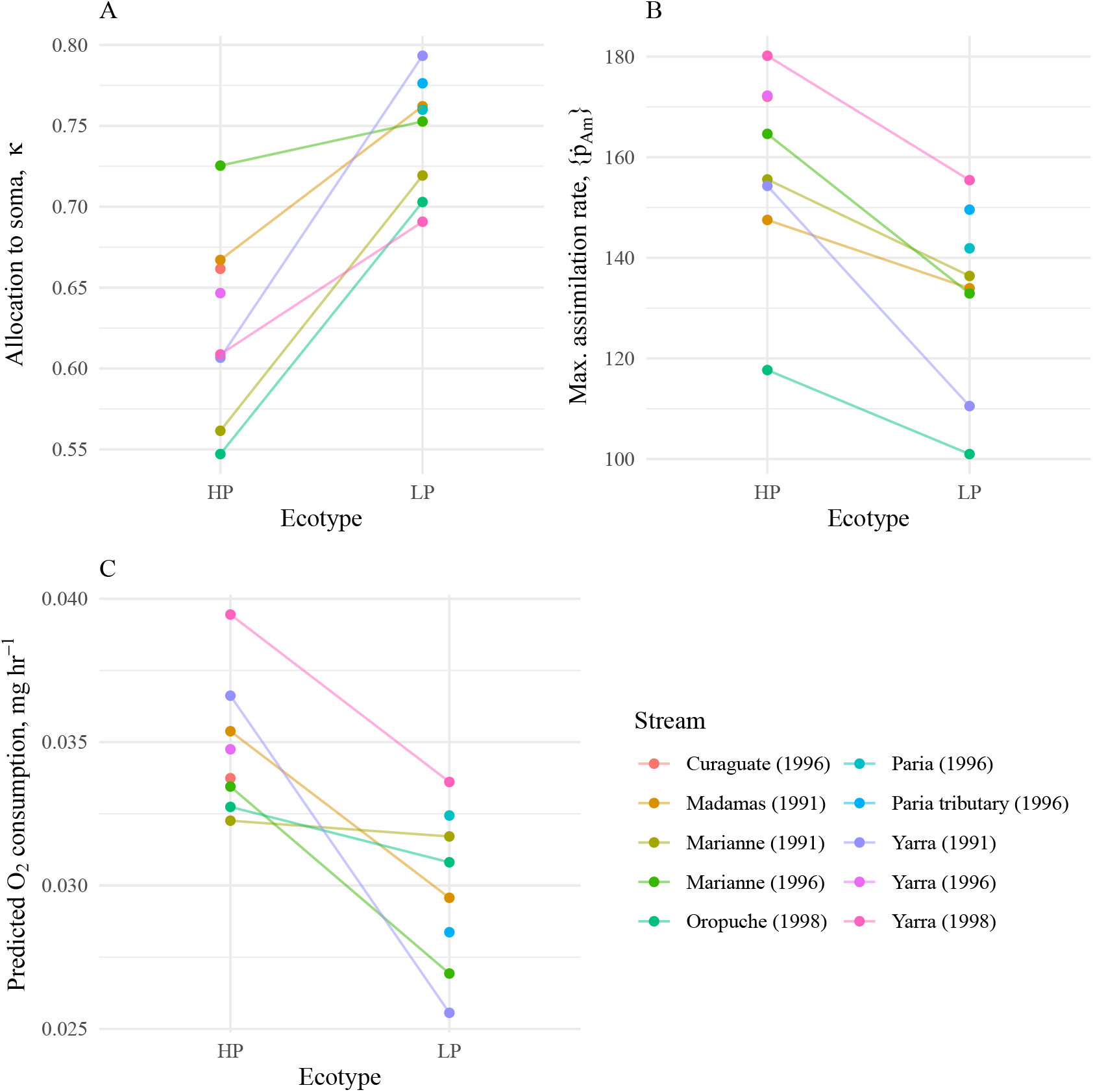
Differences between high-predation (HP) and low-predation (LP) ecotype populations in (A) the fraction of energy allocated to somatic work *κ*, (B) the maximum specific assimilation rate 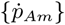, and (C) predicted basal metabolic rate (oxygen consumption), based on the full population-specific parameter set. Parameters were estimated using level 1 data availability for 16 populations. Lines connect ecotype pairs sampled from the same stream in the same year, i.e. they connect the ancestral (HP) ecotype to the derived (LP) ecotype.

### Intra- and interspecific variation

We compared the intraspecific variation in six DEB parameters reported in this study with the interspecific variation reported within the order Cyprinodontiformes (AmP, 2020) (Fig. 5). The variance in *κ* for Trinidadian guppies (across all data levels in this study) was comparable to that reported in 59 species within the order Cyprinodontiformes (*F*_23,58_ = 0.77, p=0.49). This variance in guppy *κ* values was largely an artefact due to differences in the data levels used to fit these models. When we considered only parameters from models fit with data level 1, the intraspecific variance in *κ* across 16 populations of Trinidadian guppy was significantly less than that seen within 20 species in the family Poeciliidae (*F*_15,19_ = 0.24, p=0.008). Similarly, for the five other DEB parameters (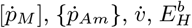, and 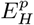), intraspecific variation was significantly less than variation within the family Poeciliidae (F-tests, p<0.05).

**Figure 5:**
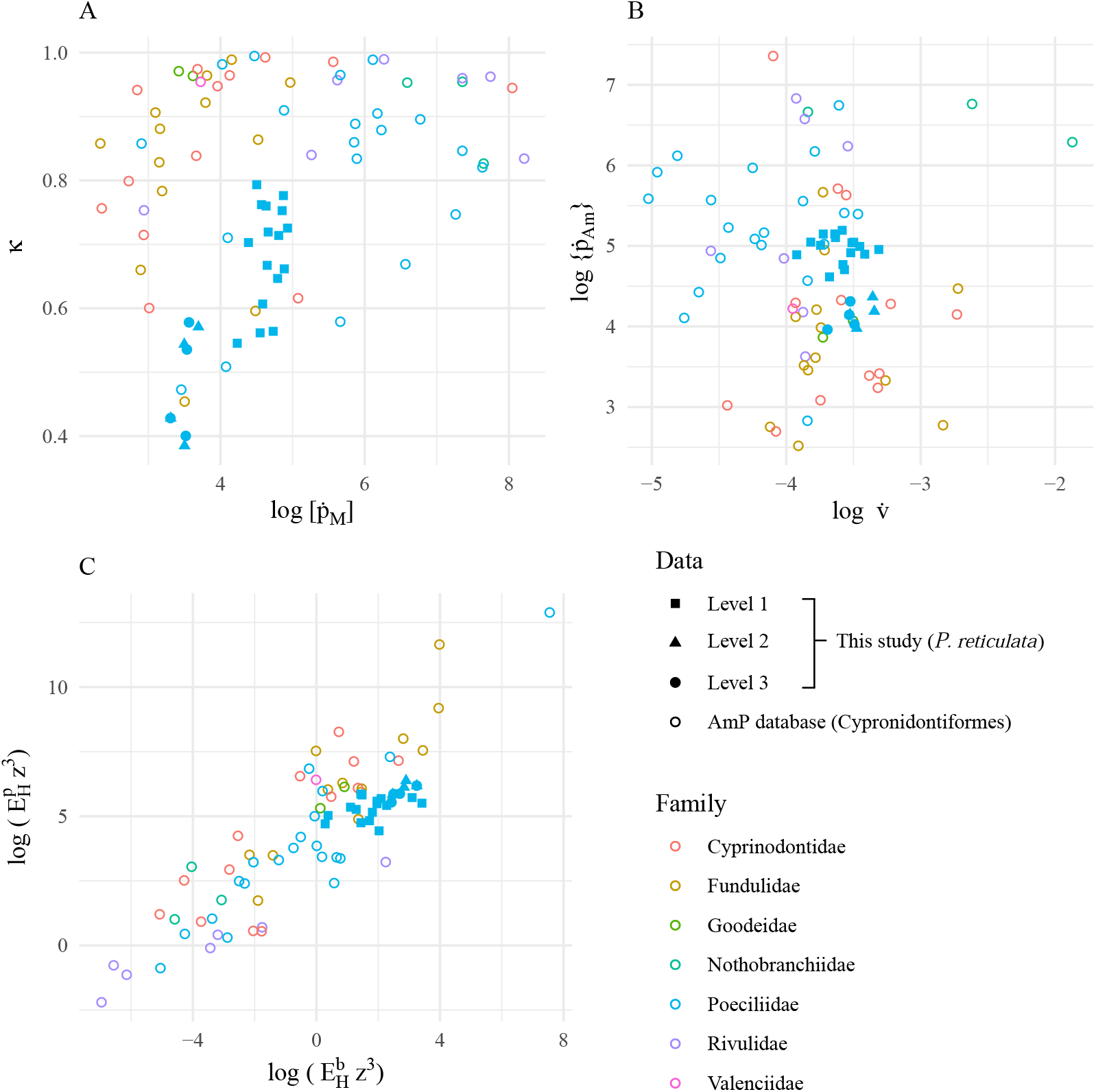
Comparison of intra- and interspecific variation in primary DEB parameters: Pairwise plots are of (A) *κ* and 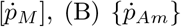 and 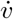, and (C) 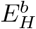 and 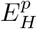. Note that to account for parameter variation due to differences in maximum body size between species, values for 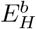 and 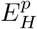 are multiplied by *z*^3^ (Kooijman, 2010). Axes are on the log-scale, except for the y-axis in (A). Filled points are parameter values obtained for Trinidadian guppies in this study, using data availability at level 1 (squares), level 2 (triangles) and level 3 (circles). For details on data levels, see Table 3. Empty circles are parameter estimates for 59 species from 7 families within the order Cyprinodontiformes, obtained from the AmP online database (AmP, 2020).

## Discussion

We identified considerable variation in dynamic energy budget (DEB) parameters among populations of Trinidadian guppies. Although variation in DEB parameters are theoretically predicted to reflect evolved, biological differences (Kooijman, 2010; Lika et al., 2011a; Marques et al., 2018; Lika et al., 2020), our results suggest that systematic bias during parameter estimation – an artefact of the type and amount of data used to fit the models – may generate substantial variation. Nevertheless, population-specific parameter sets were able to characterise known life-history variation in our study species.

### Data type influences estimation of DEB parameters

Fitting models with different types of data resulted in very different parameter estimates for the same populations. This variation was clearly an artefact of the estimation procedure, whereby the amount and type of data used to fit models had substantial effects on estimates of DEB parameters. For example, including data on metabolic rate and daily food availability halved estimates of the maximum rate at which energy is assimilated 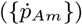; estimates of *κ* varied such that guppies in one population were predicted to allocate as much as 70% of available energy to somatic work, or as little as 40%, depending on the type of data used to fit the models (Table 4). Do these fish invest more energy in reproduction, or in somatic growth and maintenance? Had we used a single level of data availability when fitting our models, we may have felt confident in our answer. That confidence would have been misplaced: when fitting models with different types of data, our results gave conflicting answers to this basic question of energy allocation.

Nevertheless, despite these large data-type-dependent differences among parameter sets, the net combination of parameter values reproduced remarkably similar patterns of growth and reproduction, reasonably fitting the observed data (Fig 2). This is an example of getting the right answer for the wrong reasons. But which reason is least wrong? The nature of a balanced energy budget means that, for example, a decrease in somatic maintenance costs must be balanced by, say, a decrease in assimilation efficiency, if patterns of growth are to remain fixed. The interactive and parameter-rich structure of the DEB model allows many possible routes through which this balance can manifest, but those routes become increasingly constrained as more different types of data are used to fit the models.

When we included data on dioxygen consumption and daily food availability, estimates of several parameters changed substantially, and these changes were consistent across all four populations tested: maximum assimilation rate, the fraction of energy allocated to somatic work, and somatic maintenance costs all decreased, while the maturity threshold at which the adult stage is met increased (Table 4). Respectively, this meant that: less energy was entering the system from the environment; less was allocated to somatic growth; somatic overheads were reduced; and more energy was invested in attaining, and also in maintaining maturity. This happened because initial fits (without using dioxygen consumption data) generated parameter sets that overestimated metabolic rate by about a half (Table S2). When the fitting process was forced to account for the measured rates of dioxygen consumption, the total dissipation power had to decrease: data on growth and reproduction constrained this to decreases in energy assimilation rates, with more investment in non-somatic work balanced by decreases in somatic maintenance costs, and increases in the costs of maturation. As such, in the absence of dioxygen consumption data, we overestimated the energetic costs of somatic maintenance by a factor of around 2, resulting in overestimation of basal metabolic rate. We suspect that somatic and maturity maintenance costs are generally poorly determined by data on reproduction and growth alone.

### Biological variation and DEB parameters

Because the data-type-dependency of parameter estimates was consistent across the four populations that we tested, we argue that relative biological differences can be assessed when comparing DEB parameter sets fit with the same types of data. Our aim here was to determine whether the DEB approach could identify and predict known life-history variation among 16 populations of Trinidadian guppies.

Biological variation starts at the level of the individual. There were certainly biological differences among individuals in this study in their rates of assimilation and use of energy: this is apparent from the lines of data points that can be traced in Figure 2, which represent repeated measures of individuals. Although individual fish within a given food treatment received the same amount of food, there was considerable variation in maximum size and reproductive rates among individuals. Given that these fish experienced the same conditions, differences in their patterns of growth and reproduction must reflect biological differences in how each individual assimilated and allocated the energy from its food. Estimating individual-level DEB parameter sets was beyond the scope of this study; instead, we considered population-level differences in DEB parameters.

Our results demonstrate that life history variation among populations of guppies can be characterised by differences in dynamic energy budget parameters. Low-predation ecotype guppies allocated a greater fraction of energy to somatic work (higher values of *κ*), and had lower maximum assimilation rates (lower values of 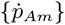) than their high-predation ecotype ancestors. These differences were consistent across 16 populations, representing five independent evolutionary origins of the low-predation ecotype (Fig. 4). For the four populations that we were able to fit at higher levels of data availability, we also found that the low-predation ecotype were less efficient at assimilating energy (lower values of *κ_X_*) than their high-predation counterparts (Fig. 3). Population-specific parameter sets gave significantly better fits to the data than parameter sets from the other ecotype, indicating that these parameter differences reflect biological differences between ecotypes.

Apart from predicting the patterns of growth and reproduction used to fit the models, how do these differences in DEB parameters compare with known life-history variation in Trinidadian guppies? Low-predation guppies invest less in reproduction (Reznick and Endler, 1982), which corresponds with the higher values of *κ* we identified here. Low-predation ecotypes also have slower metabolic rates (Auer et al., 2018)] – DEB estimates of metabolic rate from all 16 populations predicted this pattern (Fig. 4C), despite metabolic rate data not being used to fit these models.

A surprising result is lower rates and efficiencies of assimilation in low-predation guppies. Given that the low-predation ecotype evolves in response to reduced resource availability (Bassar et al., 2013; Reznick et al., 2019; Potter et al., 2021), it would appear maladaptive to evolve a reduced capacity for energy assimilation. Furthermore, low-predation guppies have increased gut length relative to the high-predation ecotype, which should facilitate higher rates of energy assimilation (Zandonà et al., 2015). However, our parameter estimates (*κ_X_* and 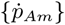) were specific to the high-protein diet used in the experiments. One possible explanation is that the experimental diet more closely resembled the natural diet of high-predation guppies, which are richer in high quality invertebrate prey (Zandonà et al., 2011). Digestion of protein-rich food is energetically costly (Cho et al., 1982), and high efficiencies may not be maintained through selection in natural low-predation populations, where diets are dominated by low-quality algae and detritus (Zandonà et al., 2011). Had a lower quality diet been used in these experiments, we may have seen higher assimilation rates and efficiencies in the low-predation ecotype.

To improve model fits, we introduced two additional parameters to the standard DEB model. First, we included a second assimilation efficiency parameter, specific to the low food treatment (*κ_X low_*). Assimilation efficiencies were consistently higher in the low food treatment than they were in the high food treatment. This could indicate that in the high food treatment not all of the daily food ration was consumed, meaning that we overestimated the amount of food eaten by individuals in this group. However, a daily record of whether each fish had consumed it’s full ration was kept for these experiments: failure to eat the full ration was rare, and there was no systematic tendency for fish in the high food treatment to not consume full rations. A more plausible explanation is that our DEB model failed to capture how total energetic flux changed as a function of food availability. One empirical measure of energetic flux - standard metabolic rate - has been shown to increase at higher food levels (Auer et al., 2015). Although this should be captured in the DEB model – reserve density should be greater, and therefore every process downstream of the assimilation flux (Table 2), should have higher rates at high food levels – in our study, this difference was partially attributed to different assimilation rates at different food levels, which is contrary to standard DEB theory (Kooijman, 2010). The second introduced parameter was the developmental delay, *t_0_*. This parameter represented a delay in the onset of development of the fertilised embryo, and was an additional term in the sub-model for age at birth. However, there is no empirical support for such a delay in guppy development (Martyn et al., 2006): the requirement of this parameter in our models suggests that energy allocation via *κ* is not constant throughout development in guppies.

Differences in parameter values between ecotypes were not always consistent between different streams. For four parameters, the magnitude of ecotype differences varied considerably between streams (Fig. 3B). Because each stream represents an independent evolutionary origin of the low-predation ecotype, these differences may reflect alternative mechanisms through which adaptation to competitive, resource-limited environments has occurred. The distinction between guppy ecotypes is well characterised (Reznick and Endler, 1982; Reznick, 1982; Reznick and Bryga, 1987). However, there is substantial life-history variation among low-predation populations from different streams, including differences in rates of senescence (Reznick et al., 2004, 2005), juvenile growth rates (Arendt and Reznick, 2005), basal metabolic rate (Auer et al., 2018), how competitive ability scales with body size (Potter et al., 2019), and in genes associated with living in a low-predation habitat (Whiting et al., 2020). Our results here support the notion that there are multiple mechanistic routes through which the low-predation ecotype can evolve.

### Implications of variation in DEB parameters

One of the strengths of the DEB approach – that models can be fit with a wide range of commonly available data – may come at a cost in terms of the reliability of comparisons of parameters across species. Although the sensitivity of parameter estimation to the types of data used to fit DEB models has been noted in a simulation study (Lika et al., 2011b), we feel that the potential scale of this problem may not be generally appreciated. Our results show that variation in DEB parameters in a single species can match the variation seen across an entire order, if differences in the types of data used to fit the models are not taken into account (Fig. 5A). This type of variation, which does not reflect true biological variation, is potentially problematic for studies that aim to quantify life-history strategies across taxa on the basis of DEB parameters (Marques et al., 2018; Lika et al., 2019; Augustine et al., 2019). However, if this type of data-type-dependent variation is truly systematic, as it appears to be in this study, then correcting for data type could improve the ability to detect such broad-scale patterns of life-history strategies.

When we considered only parameter sets fit with the same types of data, we identified intraspecific variation in DEB parameters. DEB theory predicts that closely related species should share more similar parameter values, and our results support this: guppy parameters (when fit with the same data types) tended to cluster together when compared to interspecific variation in parameters (Fig. 5), and intraspecific variation in guppies was significantly lower than the variation seen within the family Poeciliidae. However, in our system, ecotype pairs (i.e. high-predation and low-predation ecotypes) within a stream are more closely genetically related than two populations of the same ecotype, yet DEB parameters were more similar within ecotypes than between. Our results suggest that convergent evolution can result in similarity of DEB parameters that do not reflect genetic relatedness.

Our study demonstrates that variation among sets of DEB parameters can result not only from biological differences (i.e. between individuals, populations, or species), but also from artefacts of the parameter estimation process. Different parameter sets can generate the same biological patterns. In some instances this may not matter: for example, if the goal is to predict a biological process based on limited data.

In our study, DEB models predicted dioxygen consumption rates with quite impressive accuracy, given that the models were fit using data only on age, weight, length, and number of offspring. However, the goal of mechanistic modelling is to determine the causal relationships among interacting components of a system. In the context of DEB theory, this means accurately quantifying the primary parameters of the dynamic energy budget model. For DEB theory to accurately quantify the intrinsic trade-offs that underpin individual life-histories, our results suggest that data on metabolic rate and energy consumption are required when fitting dynamic energy budget models.

## Notes

### Competing Interest Statement

The authors have declared no competing interest.

### Summary of Updates

This version of the manuscript has been revised to correct some errors in the presentation of equations in Table 2. There is no change to the analyses or the results.

