## Supplementary Information for "Substantial intraspecific variation in energy budgets: biology or artefact?"

### Supplementary material for the manuscript 'Substantial intraspecific variation in energy budgets: biology or artefact?'

Tomos Potter[1*], David N. Reznick[2], Tim Coulson[1]

[1]Department of Zoology, University of Oxford, Oxford OX1 3PS
[2]Department of Evolution, Ecology and Organismal Biology, University of California, Riverside, California 92521

#### Data used to fit the models

The majority of the data used to fit population-specific DEB models came from three sets of common garden experiments (Table S1). All of these experiments followed the same general protocol, described in detail in (Reznick and Bryga, 1987). Briefly, gravid female guppies were sampled from natural high- and low-predation ecotype populations and reared for two generations under standard lab conditions. Guppies were collected in 1992, 1996, and 1998, with each year representing a different dataset: we refer to each dataset by its sampling year. Male and female guppies were crossed following a breeding plan designed to avoid inadvertent selection, and to maintain the original level of genetic variation in the wild-caught females. The data used in this study are from second-generation lab-born females. Experimental treatments started 25-30 days after the birth of the focal females. After this time, females were housed individually in 8 L aquaria and subject to either high or low levels of food. For both treatments, guppies were fed twice daily with brine shrimp and liver paste. For each fish, a daily record was kept recording whether the full ration had been consumed. There was no systematic tendency for fish in the high food treatment to not consume full rations. The exact food levels differed between the three sets of experiments (details below). Focal females were paired with mature males one week into the experiment, ensuring that they mated as soon as they attained maturity, and were re-mated following each subsequent birth. Age and standard length were recorded at the onset of the experiment, and age, standard length, and wet weight were recorded at each reproductive event. Offspring number and dry weight were recorded for each litter produced. For the 1991 and 1996 datasets, data were recorded for the first three reproductive events only. For the 1998 dataset, data were recorded at each reproductive event for the entire lifespan of experimental fish (mean age at death across all treatments and populations = 842 days). Sample sizes for each experimental population are given in Table S1.

Table S1: Population location, sampling year, and sample sizes used to estimate sixteen population-specific sets of DEB parameters. High-predation (HP) and low-predation (LP) ecotype guppies were sampled from six different river systems in three different years. Sample sizes are shown for common garden experiments in which individually housed female guppies were subject to either high or low levels of food availability. Data from fish sampled in 1991 are from (Reznick and Bryga, 1996); those sampled in 1996 are from a previously unpublished dataset (D. Reznick); data from fish sampled in 1998 are from (Reznick et al., 2004, 2005). All data and the Matlab scripts used to fit DEB models are archived on the freely-accessible online AmP database (AmP, 2020). Links to population-specific archives are provided in the table.

| River system | Sampling site name | Ecotype | Year sampled | Sample size per treatment | | Data and code archive |
|---|---|---|---|---|---|---|
| | | | | High food | Low food | |
| Oropuche | Oropuche river | HP | 1998 | 22 | 25 | tinyurl.com/y5d7a2jo |
| | Campo river | LP | 1998 | 27 | 30 | tinyurl.com/y36qq26b |
| Yarra | Yarra river | HP | 1991 | 16 | 16 | tinyurl.com/y3rc9foa |
| | | HP | 1996 | 20 | 20 | tinyurl.com/y68n2l35 |
| | | HP | 1998 | 24 | 29 | tinyurl.com/yyudxc8g |
| | Limon tributary | LP | 1991 | 15 | 15 | tinyurl.com/y5hq5dmc |
| | | LP | 1998 | 26 | 26 | tinyurl.com/y6qlykas |
| Madamas | Madamas river | HP | 1991 | 18 | 18 | tinyurl.com/y4wolu6f |
| | Miguel tributary | LP | 1991 | 17 | 17 | tinyurl.com/yy92s7to |
| Marianne | Marriane river | HP | 1991 | 15 | 15 | tinyurl.com/y5erknxb |
| | | HP | 1996 | 14 | 14 | tinyurl.com/yxsynbfm |
| | Marianne tributary | LP | 1991 | 16 | 16 | tinyurl.com/y4h5xdxt |
| | Marianito tributary | LP | 1996 | 16 | 16 | tinyurl.com/y2x97a35 |
| Curaguate | Ricon river | HP | 1996 | 16 | 16 | tinyurl.com/y6q6vd5d |
| Paria | Paria river | LP | 1996 | 15 | 15 | tinyurl.com/y6cmedu3 |
| | Paria tributary | LP | 1996 | 14 | 14 | tinyurl.com/yxb3zqdt |

High and low food treatment levels differed between datasets. For the 1998 dataset, ration size for both food treatments increased periodically over the first 5 months of the assays, and low and high food levels were designed to allow 65% and 90% of maximum growth attainable at *ad lib* food, respectively (Figure S1, (Reznick, 1980)), replicating the food availability in natural low- and high-predation habitats, respectively (Reznick, 1980, 1982). We estimated the energy content of the food rations based on calorific measurements of representative samples of both food types (Reznick, 1980). Data on the exact food levels used were not available for the 1991 and 1996 datasets, but we were able to estimate relative food treatment levels. Because the high-predation Yarra population is included in each of the three datasets (Table S1), we compared the average length at the first three partuitions across all datasets and food treatment levels. Mean standard length was greatest in the high food treatment of the 1996 dataset (mean SL=2.46 cm), indicating that this food treatment level was the highest. We then divided mean length in the other treatments by 2.46 cm to estimate relative food levels in terms of differences in patterns of growth between treatments. For the 1996 dataset, the estimated relative food abundances were 1 at high food and 0.94 at low food; for the 1991 dataset, estimated relative food abundance was 0.85 at high food, and 0.81 at low food.

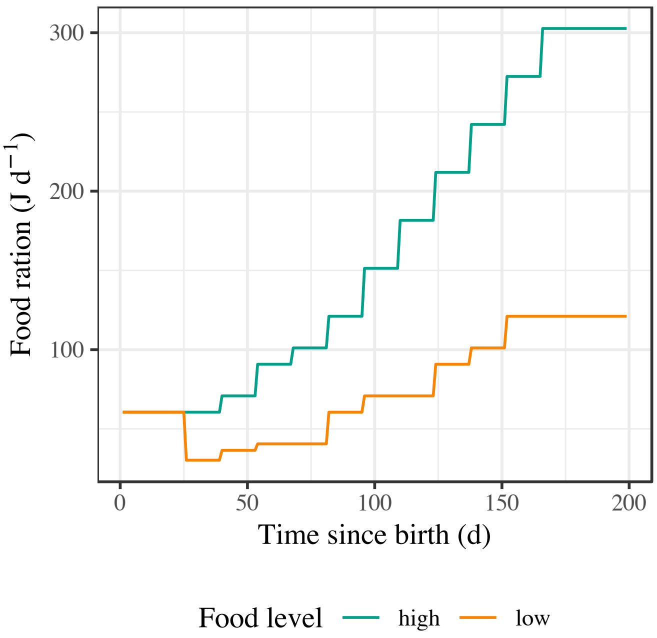


Figure S1: Quantified rations used in the 1998 experiments. 25-30 days after birth, guppies were assigned to either high (green) or low (orange) food rations. Rations increased over the first 5 months then remained constant.

The parameter estimation procedure (details in main text) requires information on maximum length at *ad lib* food levels. For the 1998 dataset, we estimated maximum length at *ad lib* food by multiplying the mean maximum length of guppies in the high-food treatment by $1/0.9$, because the high food level in this dataset was designed to allow 90% of maximum growth. Because the 1991 and 1996 datasets do not contain data on maximum size, we had to first estimate maximum growth and then adjust estimates to account for the different food levels used in these experiments. To do this, we used the 1998 dataset to fit a linear model predicting maximum length as a function of age and length at the third partuition. This model fit the data with an adjusted $R^2$ value of 0.86. We then used the parameters of this model to predict the maximum size of each individual in the 1991 and 1996 datasets, based on their sizes and ages at the third partuition. We adjusted the predicted values by multiplying each value by $1/f$, where $f$ is the relative food level in each experimental treatment (see previous paragraph). Finally, we took the mean value of our individual-level estimates, calculated for each population, as the population-specific value of maximum length at ad-lib food.

Table S2: Zero-variate observations and corresponding DEB model predictions for high-predation (HP) and low-predation (LP) ecotype guppies from the Oropuche and Yarra rivers in Trinidad. Predictions (pred) and relative error (RE) are given based on models that were fit with data availability levels 1, 2, or 3. Note that $O_2$ consumption data were not used to fit models at data levels 1 and 2, and so the respective RE terms in this table did not contribute to the overall fit.

| | | | Data level: | 1 | | 2 | | 3 | |
|---|---|---|---|---|---|---|---|---|---|
| Population | Data type | Unit | Observation | Pred | RE | Pred | RE | Pred | RE |
| Oropuche HP | Mean age at birth, high food | d | 26.01 | 26.05 | 0.002 | 25.96 | 0.002 | 25.87 | 0.006 |
| | Mean age at birth, low food | d | 26.34 | 26.30 | 0.001 | 26.40 | 0.002 | 26.69 | 0.013 |
| | Ultimate standard length | cm | 3.80 | 3.91 | 0.030 | 3.82 | 0.005 | 3.81 | 0.002 |
| | Mean dry weight at birth at high food | mg | 1.09 | - | - | 1.11 | 0.020 | 1.07 | 0.016 |
| | Mean dry weight at birth at low food | mg | 1.01 | - | - | 0.99 | 0.017 | 0.97 | 0.035 |
| | O2 consumption | mg $hr^{-1}$ | 0.025 | 0.033 | 0.315 | 0.018 | 0.280 | 0.025 | 0.015 |
| Oropuche LP | Mean age at birth, high food | d | 29.03 | 29.51 | 0.016 | 29.37 | 0.012 | 29.90 | 0.030 |
| | Mean age at birth, low food | d | 30.18 | 29.69 | 0.016 | 29.72 | 0.015 | 30.86 | 0.022 |
| | Ultimate standard length | cm | 3.51 | 3.36 | 0.044 | 3.53 | 0.004 | 3.31 | 0.057 |
| | Mean dry weight at birth at high food | mg | 1.38 | - | - | 1.49 | 0.079 | 1.36 | 0.012 |
| | Mean dry weight at birth at low food | mg | 1.45 | - | - | 1.35 | 0.072 | 1.22 | 0.157 |
| | O2 consumption | mg $hr^{-1}$ | 0.018 | 0.031 | 0.750 | 0.034 | 0.934 | 0.020 | 0.117 |
| Yarra HP | Mean age at birth, high food | d | 23.59 | 24.61 | 0.043 | 24.48 | 0.038 | 24.69 | 0.046 |
| | Mean age at birth, low food | d | 25.76 | 24.73 | 0.040 | 24.81 | 0.037 | 25.20 | 0.022 |
| | Ultimate standard length | cm | 3.59 | 3.88 | 0.080 | 3.63 | 0.012 | 3.56 | 0.009 |
| | Mean dry weight at birth at high food | mg | 0.81 | - | - | 0.85 | 0.052 | 0.81 | 0.003 |
| | Mean dry weight at birth at low food | mg | 0.81 | - | - | 0.77 | 0.054 | 0.75 | 0.078 |
| | O2 consumption | mg $hr^{-1}$ | 0.029 | 0.039 | 0.360 | 0.033 | 0.146 | 0.030 | 0.029 |
| Yarra LP | Mean age at birth, high food | d | 27.33 | 29.00 | 0.061 | 28.91 | 0.058 | 28.86 | 0.056 |
| | Mean age at birth, low food | d | 31.08 | 29.22 | 0.060 | 29.33 | 0.056 | 29.78 | 0.042 |
| | Ultimate standard length | cm | 3.33 | 3.92 | 0.175 | 3.36 | 0.008 | 3.37 | 0.011 |
| | Mean dry weight at birth at high food | mg | 1.02 | - | - | 1.12 | 0.098 | 1.10 | 0.074 |
| | Mean dry weight at birth at low food | mg | 1.10 | - | - | 1.01 | 0.087 | 0.98 | 0.105 |
| | O2 consumption | mg $hr^{-1}$ | 0.023 | 0.034 | 0.436 | 0.030 | 0.274 | 0.022 | 0.040 |

Table S3: Comparison of the mean relative error (MRE) when models are fit using the population-specific parameter set and null parameter sets. We determined the significance of the differences in fit using Wilcoxon's signed rank tests of the relative error terms of predictions to data when fit with different parameter sets.

| Population | Parameter set | MRE | Wilcoxon signed rank test | |
|---|---|---|---|---|
| | | | t | p-value |
| High Predation Oropuche | Best (HPO) | 0.076 | - | - |
| (HPO) | Null 1 (LPO) | 0.203 | 3.92 | 0.0014 |
| | Null 2 (HPY) | 0.191 | 6.71 | $7.0\times10^{-6}$ |
| High Predation Yarra | Best (HPY) | 0.086 | - | - |
| (HPY) | Null 1 (LPY) | 0.199 | 4.06 | 0.0010 |
| | Null 2 (HPO) | 0.206 | 6.71 | $7.0\times10^{-6}$ |
| Low Predation Oropuche | Best (LPO) | 0.099 | - | - |
| (LPO) | Null 1 (HPO) | 0.245 | 6.34 | $1.3\times10^{-5}$ |
| | Nul 2 (LPY) | 0.142 | 1.09 | 0.2900 |
| Low Predation Yarra | Best (LPY) | 0.085 | - | - |
| (LPY) | Null 1 (HPY) | 0.257 | 5.19 | 0.0001 |
| | Null 2 (LPO) | 0.147 | 4.06 | 0.0010 |

Table S4: Dynamic energy budget parameters for sixteen populations of Trinidadian guppy, estimated using level 1 data availability (see Table 3 in the main text for details). Parameters that were fixed, and therefore not estimated, were: the assimilation efficiency $\kappa_X$=0.25; the maximum specific searching rate $\{\dot{F}_m\} = 6.5$ L $d^{-1}$ $cm^{-2}$; the reproductive efficiency $\kappa_R = 0.95$; the surface area specific somatic maintenance $\{\dot{p}_T\} = 0$ J $d^{-1}$ $cm^{-2}$; and the Arhennius temperature = 8000 K. Parameters are given for a reference temperature of 20 °C. The mean relative error (MRE) quantifies the fit of the models to the data.

| River system | Sampling site name | Ecotype | Year sampled | Parameter: MRE | $\{\dot{p}_{Am}\}$ | $\dot{v}$ | $\kappa$ | $[\dot{p}_M]$ | $\dot{k}_{\mathcal{J}}$ | $E_G$ | $E_H^b$ | $E_H^p$ | $z$ | $\delta_M$ | $f_{high}$ | $f_{low}$ | $t_0$ |
|---|---|---|---|---|---|---|---|---|---|---|---|---|---|---|---|---|---|
| Oropuche | Oropuche river | HP | 1998 | 0.109 | 116.7 | 0.028 | 0.55 | 68.2 | 0.0019 | 5220 | 36.9 | 298.3 | 0.94 | 0.24 | 0.86 | 0.79 | 5.2 |
| | Campo river | LP | 1998 | 0.119 | 108.2 | 0.029 | 0.70 | 95.1 | 0.0020 | 5228 | 12.8 | 146.5 | 0.80 | 0.24 | 0.86 | 0.79 | 9.0 |
| Yarra | Yarra river | HP | 1991 | 0.099 | 154.3 | 0.030 | 0.61 | 98.2 | 0.0021 | 5223 | 8.2 | 306.7 | 0.95 | 0.22 | 0.85 | 0.69 | 7.0 |
| | | HP | 1996 | 0.089 | 172.2 | 0.026 | 0.65 | 120.7 | 0.0021 | 5220 | 5.4 | 446.2 | 0.92 | 0.21 | 1.00 | 0.83 | 8.0 |
| | | HP | 1998 | 0.119 | 175.0 | 0.029 | 0.61 | 120.1 | 0.0020 | 5225 | 25.2 | 382.5 | 0.89 | 0.23 | 0.86 | 0.81 | 6.0 |
| | Limon tributary | LP | 1991 | 0.118 | 110.5 | 0.028 | 0.79 | 90.4 | 0.0020 | 5220 | 4.6 | 126.2 | 0.97 | 0.24 | 0.85 | 0.71 | 8.6 |
| | | LP | 1998 | 0.137 | 156.4 | 0.027 | 0.69 | 121.0 | 0.0020 | 5222 | 8.8 | 238.9 | 0.89 | 0.23 | 0.86 | 0.76 | 9.8 |
| Madamas | Madamas river | HP | 1991 | 0.100 | 147.5 | 0.032 | 0.67 | 104.4 | 0.0020 | 5233 | 11.6 | 269.5 | 0.94 | 0.23 | 0.85 | 0.70 | 7.0 |
| | Miguel tributary | LP | 1991 | 0.134 | 134.0 | 0.033 | 0.76 | 96.5 | 0.0021 | 5191 | 6.0 | 203.6 | 1.06 | 0.24 | 0.85 | 0.67 | 7.4 |
| Marianne | Marianne river | HP | 1991 | 0.109 | 155.6 | 0.022 | 0.56 | 94.7 | 0.0022 | 5232 | 10.4 | 374.2 | 0.92 | 0.22 | 0.85 | 0.74 | 5.6 |
| | | HP | 1996 | 0.106 | 164.6 | 0.026 | 0.73 | 138.5 | 0.0020 | 5241 | 5.6 | 301.9 | 0.86 | 0.22 | 1.00 | 0.86 | 7.8 |
| | Marianne tributary | LP | 1991 | 0.123 | 136.4 | 0.030 | 0.72 | 105.9 | 0.0020 | 5216 | 7.0 | 157.0 | 0.93 | 0.24 | 0.85 | 0.67 | 7.0 |
| | Marianito tributary | LP | 1996 | 0.081 | 132.9 | 0.020 | 0.75 | 128.5 | 0.0020 | 5230 | 2.8 | 234.3 | 0.78 | 0.21 | 1.00 | 0.94 | 8.7 |
| Curaguate | Ricon river | HP | 1996 | 0.076 | 172.0 | 0.024 | 0.66 | 132.3 | 0.0021 | 5229 | 4.7 | 331.1 | 0.86 | 0.21 | 1.00 | 0.86 | 7.1 |
| Paria | Paria river | LP | 1996 | 0.101 | 141.9 | 0.036 | 0.76 | 102.9 | 0.0021 | 5219 | 3.8 | 294.0 | 1.05 | 0.25 | 1.00 | 0.83 | 8.9 |
| | Paria tributary | LP | 1996 | 0.090 | 149.6 | 0.024 | 0.78 | 130.8 | 0.0020 | 5225 | 2.1 | 218.5 | 0.89 | 0.22 | 1.00 | 0.87 | 8.5 |
